# Detoxification of host plant phenolic aglycones by the spruce budworm

**DOI:** 10.1101/472308

**Authors:** Dominic Donkor, Zahra Mirzahosseini, Jacquie Bede, Eric Bauce, Emma Despland

## Abstract

This study examines the post-ingestive fate of two host-plant derived small-molecule phenolics (the acetophenones piceol and pungenol) that have previously been shown to be toxic to the outbreaking forest pest, spruce budworm (*Choristoneura fumiferana*). We test first whether these compounds are transformed during passage through the midgut, and second whether the budworm upregulates activity of the detoxification enzyme glutathione-s-transferase (GST) in response to feeding on these compounds. Insects were reared on either foliage or artificial diet to the fourth instar, when they were transferred individually to one of two treatment diets, either control or phenolic-laced, for approximately 10 days, after which midguts were dissected out and used for Bradford soluble protein and GST enzyme activity analysis. Frass was collected and subjected to HPLC-DAD-MS. HPLC showed that the acetophenones do not autoxidize under midgut pH conditions, but that glucose- and glutathione-conjugates are present in the frass of insects fed the phenolic-laced diet. GST enzyme activity increases in insects fed the phenolic-laced diet, in both neutral pH and alkaline assays. These data show that the spruce budwom exhibits counter-adaptations to plant phenolics similar to those seen in angiosperm feeders, upregulating an important detoxifying enzyme (GST) and partially conjugating these acetophenones prior to elimination, but that these counter-measures are not totally effective at mitigating toxic effects of the ingested compounds in the context of our artifical-diet based laboratory experiment.

## Introduction

In the chemical arms race between plants and herbivores, insects exhibit multiple mechanisms for dealing with plant secondary metabolites in their diet, ranging from avoidance, rapid egestion, enzymatic detoxification via degradation or conjugation, or even sequestration for use in defense [1]. Insect counter-defenses are often phenotypically plastic, especially in generalist feeders, and induced by plant compounds in the diet. These counter-adaptations to plant defenses can also occur at different time points: before ingestion, in the digestive tract prior to absorption or within cells afterwards [2,3].

The midgut is the main site of digestion in insects and a major interface for detoxification of plant allelochemicals [4]. The midgut is lined by a peritrophic membrane, a sheath of chitin microfibrils crosslinked by proteins around the midgut lumen that prevents large molecules from entering midgut cells, and thus constitutes a first line of defense against absorption of allelochemicals [5]. In many Lepidopteran species, the midgut lumen enclosed by the peritrophic membrane is highly alkaline, which in itself can be considered another counter-adaptation to plant defenses, since high pH decreases the protein binding capacity of tannins, improves the extraction of protein from leaf tissue and contributes to inactivating plant defensive enzymes [6,7].

Phenolics are one of the most important classes of plant secondary metabolites. Their detrimental effects have been linked to oxidation of biologically important molecules such as proteins and DNA. Large-molecule phenolics such as tannins can be either autoxidized or oxidized by plant enzymes in the midgut lumen, generating low molecular weight ROS that cross the peritrophic membrane to enter cells [8,9]. However, smaller phenolic compounds could cross the peritrophic membrane themselves and cause lesions and oxidative stress in cells directly [5]. Oxidative stress in midgut tissues, caused by either of the above mechanisms, is associated with lower caterpillar performance [8].

One approach to understanding insect counter-adaptations to phenolic compounds involves following the fate of phenolics in Lepidopteran midguts by assaying the original compounds and their metabolites in the insects’ frass. These studies suggest that the outcome varies greatly depending on the compounds’ structure: some move through the digestive tract of the caterpillar intact, while others are modified by enzymes and eliminated in a less toxic form [10]. Enzymatic detoxification depends on the compound and insect species, and can involve glycosylation, glutathionation, sulfation or deacylation [10-13]. A broad range of detoxification enzymes have been recorded from insect midguts [14]: many are located in the cytoplasm of midgut cells where they act to prevent damage to biological molecules and hasten excretion of toxic compounds, but some are secreted into the midgut lumen where they act on plant toxins before they enter cells [3]. Lepidopteran digestive enzymes are known to be adapted to function in alkaline conditions [15], and the same is likely to apply to secreted detoxification enzymes [14].

Most work on phenolic mechanisms of toxicity and insect counter-adaptations has been with angiosperm feeders [8]. However, phenolic compounds are known to play a key role in conifer defense against herbivores [16], and the composition of conifer phenolics differs from those found in angiosperms [9].

Two phenolic compounds have been suggested to play an important role in the defense of coniferous trees against defoliation by the spruce budworm, *Choristoneura fumiferana* Clemens (Lepidoptera: Tortricidae) [17,18], the most serious insect pest of coniferous forests of eastern North America [19]. Two sets of acetophenones have been identified from white spruce (*Picea glauca*) trees resistant to budworm attack which suffered only light defoliation when other trees around them were heavily damaged [20]: piceol and pungenol (aglycones) and picein and pungenin (their glycosides). The aglycones increase mortality and slow growth in bioassays, but the glycosylated forms appear to have no effect on the budworm [18].

These findings have turned a spotlight onto the role of acetophenones in conifer defense against folivores and in particular, against the spruce budworm. Since then, these acetophenones have been shown to be broadly distributed across coniferous trees: ten of 12 surveyed Pinaceae species accumulated at least one of the two glycosylated acetophenones in the foliage, of which 4 species accumulated both a glycoside and the corresponding aglycon. In general, the glycosides were found alone or at higher concentrations than the aglycones [21]. Genetic analysis of white spruce trees showed that a glucosyl hydrolase gene, *PgBgluc-1*, was constitutively highly expressed in resistant trees, catalyzing formation of the aglycones from the glycosylated compounds [22]. Levels of both the gene transcripts and the aglycones are highly heritable and are thought to be maintained via selection pressures imposed by spruce budworm herbivory [17].

As their name implies, spruce budworm are early-spring feeders, attacking buds as they begin to elongate. In white spruce, the aglycones begin to be expressed in current-year foliage near the end of shoot elongation, when the budworm reach the final instars [17]. Budworm feeding on white spruce are therefore exposed to these compounds in the final larval instars, as well as early in the season when insects emerging from diapause feed on previous-year foliage before buds become available. The final-instar spruce budworm midgut has a highly alkaline pH (10.5 ± 0.12 [23]) and has been shown to express at least one major detoxification enzyme, glutathione-s-transferase (GST) in response to feeding on foliage [24]. The present study aims to determine the fate of piceol and pungenol after ingestion by the budworm, testing first whether these compounds are transformed during passage through the midgut or eliminated unchanged, and second whether the budworm upregulates GST activity in response to feeding on these compounds.

## Materials and Methods

### Experimental design

Spruce budworm insects were obtained at the second instar larval diapausing stage from the Great Lakes Forest Research Centre, (Canadian Forest Service, Sault Ste. Marie, ON, Canada). The larvae were placed in groups of 10 in Solo cups (2 cm diameter, 4 cm long) and reared in a laboratory incubator on pre-treatment diet (either white spruce foliage or artificial diet) at 23°C, 50% relative humidity. The two pre-treatment diets (foliage or artificial diet) were used to control for possible down-regulation of detoxification enzymes when feeding on artificial diet rather than foliage [24]. The experiment was run 3 times (once on foliage in 2016, twice on artificial diet (2016 & 2017)).

At the moult to the fourth instar, larvae were placed individually in new cups containing the treatment diet (either control or phenolic-laced) until one week after the moult to the sixth instar when they were removed for immediate use in the experiment (N=48 larvae per treatment per experimental run).

The experiment began by weighing the caterpillars, then dissecting them to remove midguts for immediate biochemical analyses, specifically Bradford soluble proteins and glutathione-*S*-transferase activity. Frass from the treatment cups was collected and frozen at - 80°C until HPLC analysis.

### Insect diets

In the foliage pre-treatment, insects were reared until fourth instar on fresh current-year white spruce foliage collected at Morgan Arboretum (45°53’N, 72°92’W): twigs were placed in water picks and changed every three days to ensure freshness. In the artificial diet pre-treatment, initial rearing was done on modified McMorran Grisdale artificial diet [25] prepared in the laboratory as per the recipe provided by Insect Production Services, Canadian Forest Service (Sault Ste. Marie, ON, Canada).

Piceol (4’-hydroxy-acetophenone) and pungenol (3’, 4’-dihydroxy-acetophenone) were purchased from Sigma-Aldrich (Oakville, ON, Canada). To prepare the phenolic-laced diets, 0.966 g each of the two compounds (to achieve the upper range of physiological concentrations [18]) was dissolved in 10 ml of methanol and added to the artificial diet at the same time as the vitamin mixture.

### HPLC-DAD-MS analysis

High Performance Liquid Chromatography-Diode Array Detector-Mass Spectroscopy (HPLC-DAD-MS) was used for chromatographic separation and identification of phenolics in the budworm frass.

Since it was suspected that the alkaline environment of the caterpillar midgut may lead to oxidative reactions [26], a mix of piceol and pungenol (1 mg/ml each) was incubated at either neutral (pH 7.2) or alkaline (pH 9.5) conditions for 24 hours and subjected to HPLC-DAD-MS.

Phenolic compounds were extracted from the frass of spruce budworm pre-treated on artificial diet (2017 run) following [22]. Frass from 30 individual caterpillars from each treatment diet was pooled and dried in an oven for 24 hrs and ground to powdered form using liquid nitrogen, then stored in 2 ml Eppendorf tubes at −80°C prior to analysis. 50–100 mg of fine dried frass powder was extracted using 1 ml of 70% HPLC grade methanol. Benzoic acid (1 mg/ml) was used as an internal standard with 150 μl of benzoic acid added to 350 μl of the liquid sample. 70% methanol (600 μl) was added to the frass powder and incubated at 4°C on a shaker. After 6, 24 and 48 hours of incubation, the samples were centrifuged at 13 000 g for 10 mins. The supernatants were pooled and kept at −80°C. A fresh 600 μl of aqueous methanol was added to each sample, and after incubation, centrifugation was repeated. Extracts obtained after 6, 24 or 48 hours were pooled as a single extract for HPLC-DAD-MS analyses. Extraction and analysis was replicated three times.

Phenolics were separated by HPLC-DAD-MS using a Spursil C18 3 μm column (150 x 2.1 mm) maintained at 25°C. The mobile phase consisted of (A) 0.1% formic acid and (B) 0.1% formic acid in acetonitrile (ACN). The separation gradient was as follows: 0-12 min, 3-45% B; 12-13 min, 45-95% B; 13-15 min, 95% B and 15-18 min, 95-98% B. The column flow rate was 250 μl per min. The detection wavelength was at 280 nm. For mass spectrometry, a micromass quantitative-time-of-flight (q-Tof) spectrometer (Ultima ^TM^ API instrument) with electrospray ionization (ESI) in the positive mode was used for detection and identification of conjugated forms of the phenolic compounds. The following parameters were used: scanning range between *m/z* 200-500, 3.5 K volt with a scan time of 1 second, drying gas flow 6 mL/min, nebulizer pressure 60 psi, dry gas temperature 300 °C, vaporizer temperature 250 °C. The instrument was programmed to detect compounds with a molecular mass between 50 to 900 Da.

### Biochemical analyses

In each of the three runs of the experiment, sixth instar caterpillars were dissected to remove the midguts. Four midguts were pooled for each sample. Midguts were rinsed in saline dissection buffer and placed four together in an Eppendorf tube that contained 600 μl of sterile dissection buffer and 600 μl of 2X protease inhibitor cocktail. The midgut samples were homogenized and the homogenates centrifuged at 13 000 rpm at 4°C for 5 minutes. The supernatants were transferred to new, sterile Eppendorf tubes. 5 μl aliquots of gut homogenate were used for the soluble protein assay and 10 μl aliquots of the homogenate was used for the assay measuring GST activity on the same day as dissection. Enzyme assays were conducted at both neutral and alkaline pH, to mimic conditions inside cells and in the midgut lumen [23].

### Soluble protein assay

The soluble protein concentration of each midgut sample was determined by the use of the modified Bradford assay (Bradford, 1976; Zor & Ernst, 2010). The buffer used was 0.1M phosphate buffer, pH 7.2. The Bradford reagent (Bio-Rad) was diluted in distilled water as per manufacturer’s instructions. The linear concentration range was between 0.1-1.4 mg/ml. Bovine serum albumin (BSA) was used to make a standard curve. All samples and standards were analyzed in triplicate and a negative control was included. The absorbance was measured at both 590 nm and 450 nm using an Infinite PRO 200 spectrophotometer (Tecan). The soluble protein concentration of the unknown samples was determined by comparing the A_590/450_ values against the standard curve.

### Glutathione-S-transferase enzyme assay

Glutathione-S-transferase (GST) catalyzes the addition of reduced glutathione (GSH) to the substrate, in this case 1-chloro 2, 4 dinitrobenzene (CDNB). The product of the reaction formed is a yellow colored product that can be monitored at 340 nm. Each 96-well plate included a positive control (GSH, buffer, GST enzyme and CDNB), a negative control (GSH, CDNB and buffer) and experimental samples (GSH, CDNB, gut homogenate and buffer), all of which were assayed in triplicate. The assay was conducted at both neutral pH, 100 mM potassium bicarbonate buffer (pH 7.2) as well as at alkaline pH, 100 mM sodium bicarbonate buffer (pH 9.2). Each reaction mixture contained the following: 25 μl of 10 mM GSH, 25 μl 10 mM CDNB dissolved in 0.1% v/v in 95% ethanol, 10 μl of 0.1 U/ml GST enzyme, 10 μl of gut homogenate, and was transferred into a ultraviolet microplate well that contained appropriate buffer up to a total volume of 250 μL. Enzyme activity was determined by monitoring changes in absorbance at 340 nm measured every 15 seconds for 2 minutes under the spectrophotometric kinetic mode, at a constant temperature of 25°C.

### Statistical analysis

Caterpillar mass at the beginning of the experiment was analyzed by two-factor Anova, in order to evaluate the effects of the acetophenones on growth, as a proxy for performance. Soluble protein and GST activity were compared between the phenolic-laced and control diets with separate t-tests for each run of the experiment: these data could not be compared directly between different runs of the experiment as assays were conducted separately.

## Results

### Caterpillar mass

Two-way analysis of variance showed that growth of budworm was lower on phenolic than on control diet (F_1, 269_ = 7.62, p = 0.006). Growth also differed between the three runs of the experiment (F_2, 269_ = 111.9, p <0.001), being lower on the artificial diet than on the foliage pre-treatment, see Figure 1. The interaction term was not significant (F_2, 269_ = 1.44, p = 0.23).

**Figure 1:**
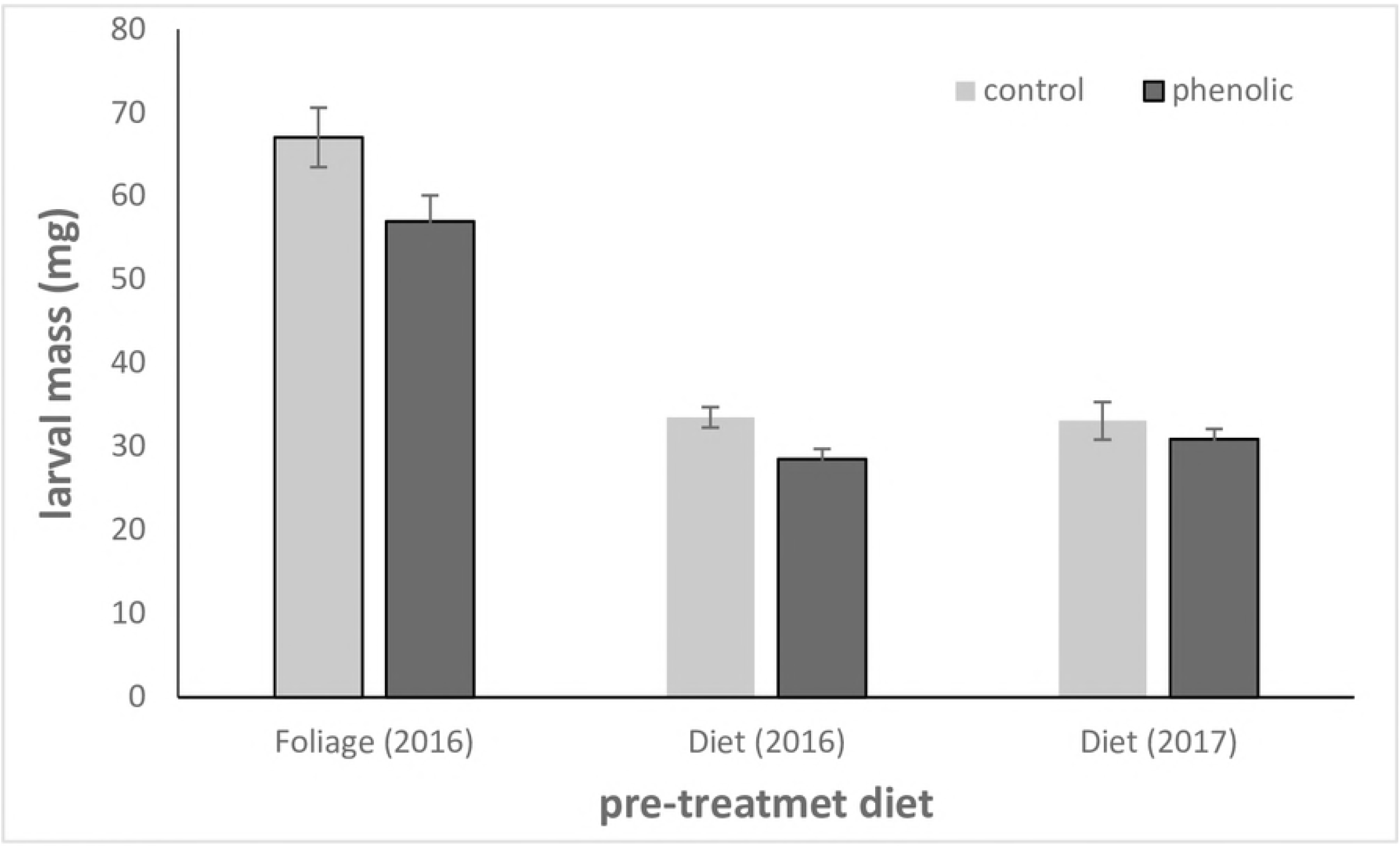
Body mass of sixth instar budworm caterpillars (mean±SE, N=50) fed on either control- or phenolic-laced artificial diet in the three experimental runs (pretreatment on foliage (2016) or on artificial diet (2016 & 2017)).

### HPLC-DAD-MS of incubated phenolic solutions

Peaks corresponding to piceol and pungenol standards were observed in the mixed solutions incubated at neutral and alkaline pH. Several other peaks were also observed after incubation in the alkaline solution (Figure 2.) One of these peaks was also observed after incubation at neutral pH at a similar level, but the other two were at much lower concentrations. The molecular masses (m/z 285, 287 and 303) of these compounds suggest that they may be dimers of piceol (m/z 136.45) and pungenol (m/z 152.15).

**Figure 2:**
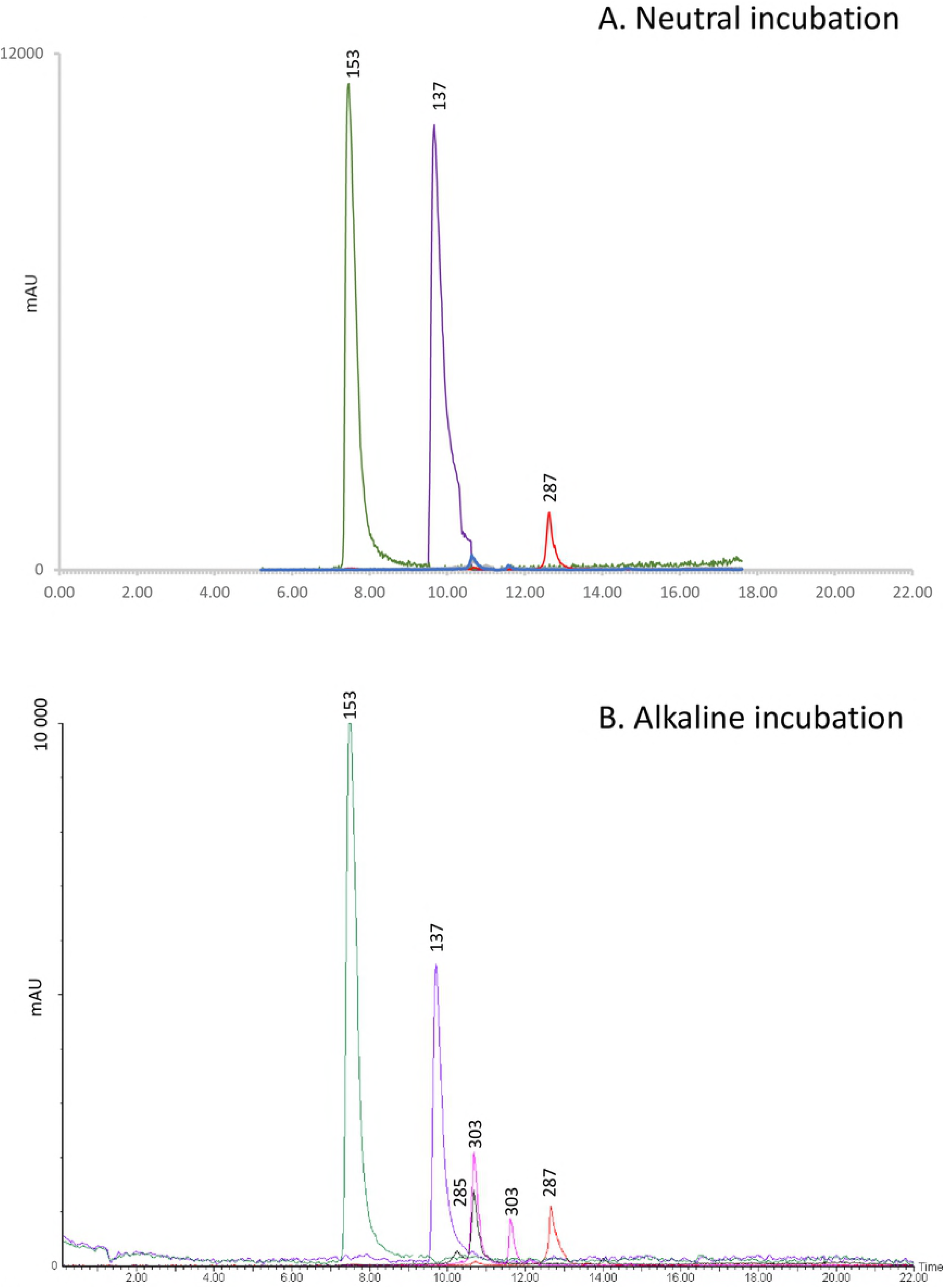
HPLC-DAD chromatograms of the piceol/pungenol mixture incubated at A. neutral and B. alkaline pH. Numbers represent m/z of identified peaks: 137 = piceol, 153 = pungenol, 285, 287 and 303 = putative dimers.

### HPLC-DAD-MS of budworm frass

In the frass from caterpillars fed on phenolic diet, peaks were detected corresponding to piceol (m/z 136.45) and pungenol (m/z 152.15) and to one of the putative dimers observed in the alkaline incubation (m/z 287) (Figure 3). Four additional peaks were also observed, and tentatively identified as the phenolic glycosides picein (m/z 299.09) and pungenin (m/z 315.11), and as the glutathionylated conjugates of piceol (m/z 425.28) and pungenol (m/z 441.23). These m/z represent the loss of water plus addition of glutathione to the phenolic and the elimination of a proton. As expected, frass from caterpillars fed on control diet did not contain any of these compounds.

**Figure 3:**
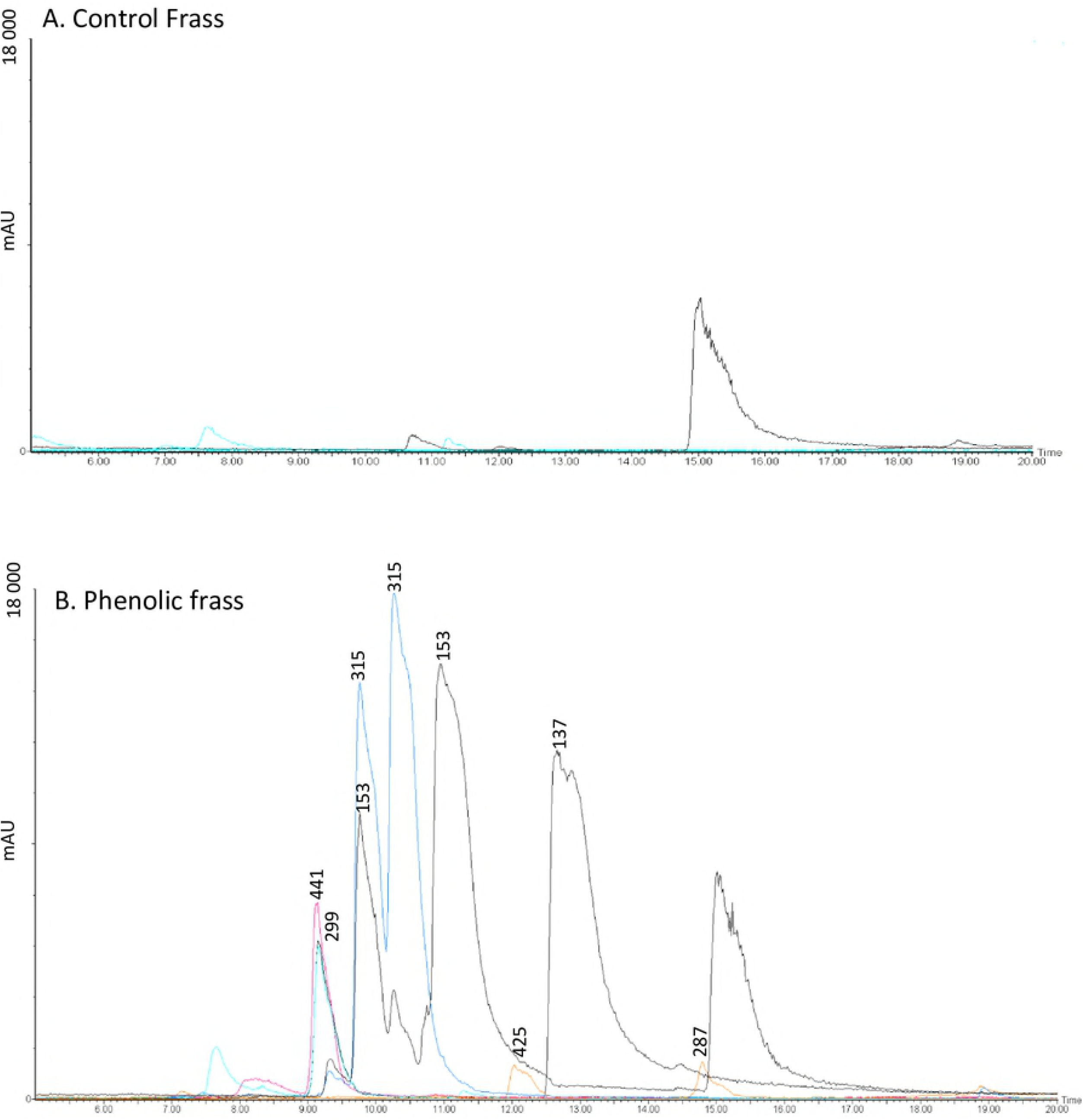
HPLC-DAD chromatograms of frass from caterpillars fed A. control or B. phenolic-laced artificial diet. Numbers indicate m/z of identified peaks: 137 = piceol, 153 = pungenol, 287 = putative dimer, 299 = tentatively identified as picein, 315 = tentatively identified as pungenin, 425 = tentatively identified as a GSH-piceol conjugate, 441 = tentatively identified as a GSH-pungenol conjugate.

### Midgut soluble protein levels

The midgut soluble protein levels of the spruce budworm caterpillars fed on control diet were not significantly different from those of budworm fed on phenolic diet in any of the three experimental trials: foliage to artificial diet: t_22_ = −1.52, p = 0.14; artificial to artificial diet: t_22_ = - 1.59, p = 0.14 (2016); t_22_ = −0.622, p = 0.54 (2017).

### Glutathione-S-transferase activity

At alkaline pH, glutathione-*S*-transferase enzyme activity in the midgut of the budworm fed on phenolic diet was significantly higher than in controls, in the foliage-to-artificial-diet experiment (t_22_ = −4.03, p = 0.0006), as well as in the artificial-to-artificial-diet experiment: t_22_ = −3.20, p = 0.004 (2016) & t_22_ = −2.64, p = 0.017 (2017). See Figure 4 **Error! Reference source not found.**. At neutral pH, GST activity also increased on phenolic relative to control diet: foliage to artificial diet: t_22_ = −2.11, p = 0.05; artificial to artificial diet: t_22_ = −4.28, p = 0.0007 (2016); t_22_ = −3.53, p = 0.004 (2017). See Figure 4 **Error! Reference source not found.**.

**Figure 4:**
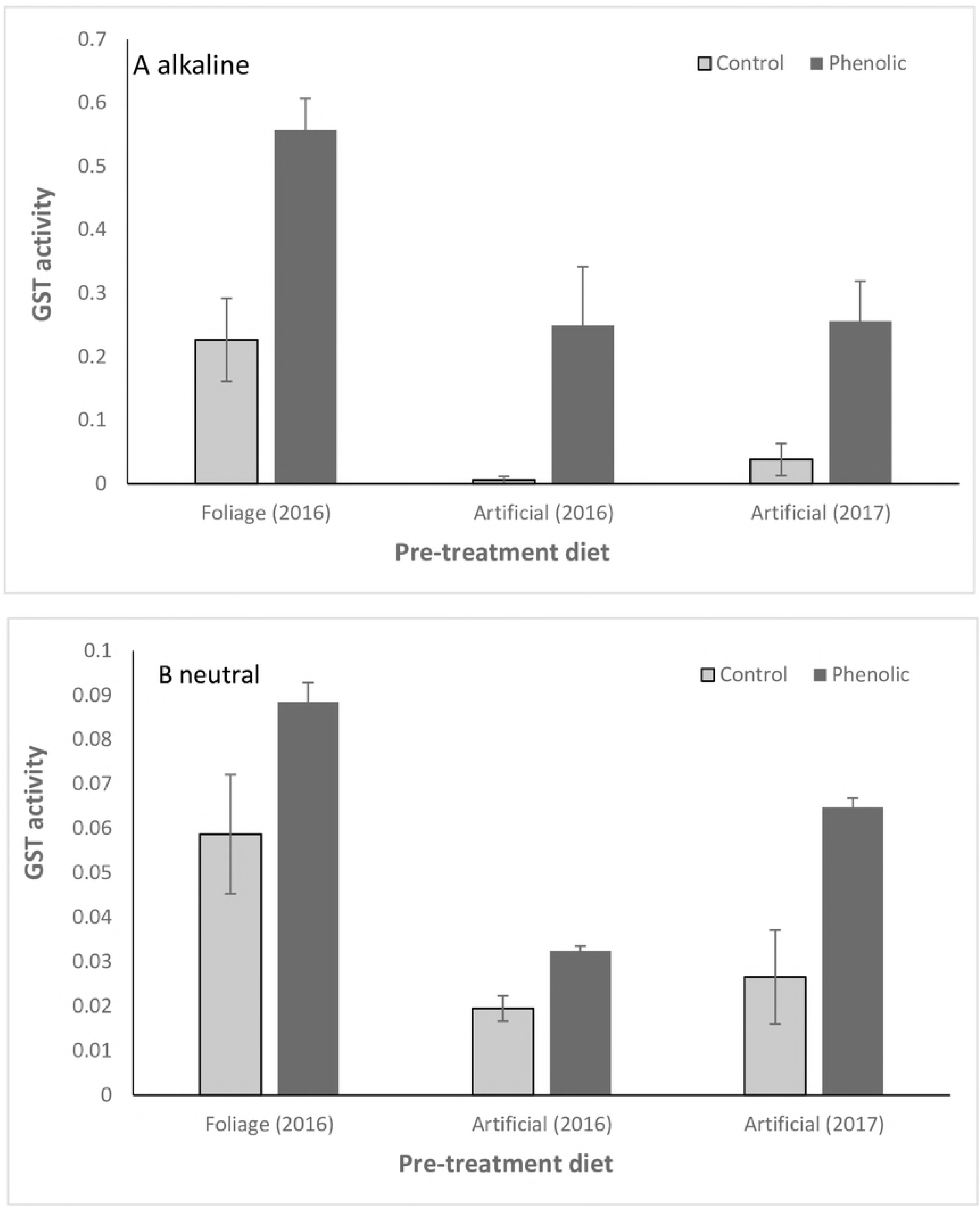
Glutathione-S-transferase enzyme activity of spruce budwormthe midgut tissue of the spruce budworm at neutral and alkaline pH, represented per mg soluble protein (mean ± SE, N=12).

## Discussion

Comparison of HPLC-DAD-MS chromatograms of acetophenones incubated at neutral and alkaline pH suggests that these compounds do not autoxidize under alkaline conditions. Similarly, previous work suggests that the phenolics of *Picea abies*, unlike those of many angiosperms, show only minor changes under alkaline incubation [26]. Interestingly, the chromatograms suggest that the acetophenones partially dimerize under high pH, but the biological significance of this effect is not clear.

HPLC-DAD-MS chromatograms of frass of insects fed acetophenone-laced diet showed, in addition to the original compounds and their dimers, novel peaks not observed in the alkaline-incubated solutions. Based on m/z, these compounds are putatively identified as glycosylated and glutathionated conjugates of the original piceol and pungenol. As expected, none of these compounds were detected in the frass of control insects. The novel peaks in the phenolic frass are therefore likely to be the products of enzymatic metabolism. Indeed, our results further show upregulation of glutathione-S-transferase in insects fed phenolic-laced artificial diet compared to controls.

Glycosylation and glutathionation of the acetophenones likely represent detoxification prior to elimination [10]. The acetophenone glycosides (picein and pungenin) are known to be less harmful to the budworm than are the aglycones [18]. Indeed, glycosides are often less toxic than their aglycones [2], and glycosylation is a common form of detoxification [3]. For instance, in *Epirrita autumnata*, phenolic glycosides were egested without metabolic modifications, but that phenolic aglycones were glycosylated prior to egestion{{23291 Salminen, J.P. 2004}}. The enzyme family likely responsible for glycosylation are the UDP-glycosyltransferases (UGTs) whose role in insect detoxification of plant compounds is as yet poorly understood [3]. UGTs have been demonstrated in several folivorous Lepidoptera [3,27,28], but not the spruce budworm; they have been demonstrated in white spruce and linked to glycosylation of the acetophenone aglycones for storage [29].

Glutathionation is better understood as a detoxification mechanism used by herbivorous insects, and detoxification of plant chemicals and/or insecticides via GST activitiy has been studied in numerous Lepidoptera [30], including the spruce budworm [24]. GSTs are a complex and widespread enzyme superfamily: multiple GSTs have been detected in all insects studied to-date, with variable specificities for a range of compounds. Generally insect GSTs catalyse the conjugation of reduced glutathione (GSH) to electrophilic molecules; they thus generate glutathione-S-conjugates that are more water soluble {{23689 Enayati, A.A. 2005}}, and can prevent oxidative stress [31]. GSTs can be upregulated in response to plant allelochemicals in the diet [30], and glutathione conjugates of plant metabolites have been detected in the frass of several caterpillar species [12].

Previous work on the spruce budworm has shown higher levels of expression of *Choristoneura fumiferana* GST mRNA and proteins in whole body extracts of sixth instar larvae fed on balsam fir (*Abies balsamea*) foliage compared to caterpillars reared on artificial diet [24]. The authors suggested that GST could play an important role in detoxifying host secondary metabolites, but could not identify the compounds responsible. Our results demonstrate that GST is upregulated in the midgut in response to harmful aglycone acetophenones, and that these compounds are partially glutathionated prior to elimination. Comparison of GST activity between the two pre-treatment diets (foliage vs artificial diet) also suggests upregulation of the enzyme prior to the beginning of the experiment in response to compounds in the white spruce foliage [24].

Detoxification via GST activity has been recorded in the insect fat body, midgut and haemolymph [30,32,33]. Most studies record intracellular GST activity as contributing to phase II detoxification by binding lipophilic compounds so they are more easily removed from the cell [32]. However, it remains unclear whether GST could also be secreted into the midgut lumen. GST-catalysed conjugation neutralizes the electrophilic sites of lipophilic substrates by attaching reduced GSH. GSTs have a high affinity towards GSH, which is generally present at high concentrations intracellularly [12]. However, GSH has also been detected in caterpillar midguts, and the ratio of reduced to oxidized GSH in the midgut lumen is used as an index of oxidative stress from plant phenolics [34,35]. Schramm{{23662 Schramm, K. 2011}} detected GSH conjugation to plant allelochemicals at high pH conditions representing those of caterpillar midguts, but were unable to disentangle non-enzymatic from GST-catalyzed reactions. We detected higher GST activity at alkaline than at neutral pH, suggesting that spruce budworm GSTs could function in the midgut lumen. Previous studies suggest that GSH in the midgut lumen could come, at least in part, from foliage, and spruce needles have been shown to contain GSH [36]. Detoxification of plant compounds by enzymes in midgut tissues has been much studied [3], but non-absorption of defensive chemicals due to conjugation in the midgut lumen warrants further attention [14], since it would be a very powerful adaptive counter-measure to neutralize toxins before they even contact damageable tissues.

Low molecular weight phenolics from angiosperms have been shown to cause oxidative stress in caterpillar midgut tissues but it is not always clear whether the damage is caused by ROS created by oxidation of the phenolics in the midgut lumen or directly by the phenolics themselves after their absorption in midgut tissues {{72 Barbehenn,R V. 2011}}. We show no evidence of oxidation of piceol and pungenol under high pH conditions, consistent with previous work on *Picea* phenolics. If small compounds like acetophenones can cross the peritrophic membrane and directly damage tissues, secreted GSTs could limit absorption of these compounds by making them larger and more hydrophilic and less likely to be absorbed, and cytosolic GSTs could limit damage caused by phenolics that have crossed the peritrophic membrane and been absorbed. We show GST activity at both neutral and alkaline pH, but our results cannot resolve whether the enzymes responsible are cytosolic or secreted. In general, few studies pinpoint the exact location of enzyme activity [14], leaving it a question open to further investigation.

Finally, our results showed that the acetophenones are only partially conjugated and that they still negatively affect budworm performance, at least in the context of an artificial diet-based laboratory assay. Indeed, addition of piceol and pungenol to artificial diet reduced growth of spruce budworm larvae in comparison to caterpillars fed control artificial diet. Similarly, Schramm {{23662 Schramm, K. 2011}} concluded that GST-conjugation of ingested plant compounds occurs, but only partially, and is of limited importance relative to other detoxification mechanisms, including rapid elimination. One possible explanation is that this conjugation is costly, as GSH contains 3 Nitrogen atoms, a nutrient often limiting in herbivores, including the spruce budworm [37]. The sulfur amino acids required for GSH synthesis are particularly low in foliage: for instance, close to 20% of sulfur amino acids ingested by two angiosperm-feeding caterpillars are directed to GSH production [38]. This would support a well-known relationship between protein nutrition and allelochemicals, whereby insects deficient in dietary N are often more sensitive to plant defensive compounds [39].

In general, organisms with an evolutionary history of consuming phenolics are less sensitive to their deleterious effects, via physical and biochemical defenses in the midgut including surfactants, high pH, low redox potential, low oxygen levels, antioxidants, antioxidant enzymes, and detoxification enzymes [8]. We show that the spruce budworm exhibits counter-adaptations to plant phenolics similar to those seen in angiosperm feeders, upregulating an important detoxifying enzyme (GST) and partially conjugating these acetophenones prior to elimination, but that these counter-measures are not totally effective at neutralizing effects of the ingested compounds in the context of our artificial-diet based laboratory experiment. As a further twist in the co-evolutionary arms race between plants and their insect herbivores, conifers have been shown to produce flavonoids that act as GST inhibitors in vitro and increase insecticide mortality; these could thus represent counter-counter-adaptations against insect detoxifying enzymes that synergize the plants’ defensive phenolics [40].

## Acknowledgements

Thanks to the insect production services of the Canadian Forest Service for providing the insects. Thanks to Alain Tessier of Concordia University’s Center for the Biological Applications of Mass Spectroscopy for help with the HPLC. This work was funded by an NSERC Discovery grant to E.D. (RGPIN/250173-2002), a FQRNT Aménagement et Environnement Forestiers V grant to E.D. and E.B. (186342) and by a Concordia University Dean’s Award of Excellence to D.D.

